# Epigenetic bookmarking of H_2_S exposure in *Caenorhabditis elegans*

**DOI:** 10.1101/619734

**Authors:** Emily M. Fawcett, Jenna K. Johnson, Chris Braden, Esteli M. Garcia, Dana L. Miller

## Abstract

Physiological memories of environmental stress can serve to predict future environmental changes, allowing the organism to initiate protective mechanisms and survive. Although physiological memories, or bookmarks, of environmental stress have been described in a wide range of organisms, from bacteria to plants to humans, the mechanism by which these memories persist in the absence of stress is still largely unknown. We have discovered that *C. elegans* transiently exposed to low doses of hydrogen sulfide (H_2_S) survive subsequent exposure to otherwise lethal H_2_S concentrations and induce H_2_S-responsive transcripts more robustly than naïve controls. H_2_S bookmarking can occur at any developmental stage and persists through cell divisions and development but is erased by fasting. We show that maintenance of the H_2_S bookmark requires the SET-2 histone methyltransferase and the CoREST-like demethylase complex. We propose a model in which exposure to low doses of H_2_S generates a long-lasting, epigenetic memory by modulating H3K4me2 modifications at specific promoters. Understanding the fundamental aspects of H_2_S bookmarking in this tractable system can provide mechanistic insight into how environmental exposures are translated into the epigenetic landscape in animals.

## Introduction

One strategy animals use to maximize survival in a changing environment is to anticipate changes based on prior experiences. In some situations, physiological adaptations to environmental changes persist long after the initial stimulus has been removed. These sustained cellular memories of previous life experiences are known as environmental bookmarks (reviewed in (Kinoshita and Seki, 2014). Perhaps the best-studied example is in the budding yeast *Saccharomyces cerevisiae*, where cells which have previously been grown in media containing galactose respond to subsequent galactose exposure faster than naïve yeast (Kundu et al., 2007). Similarly, human retinal cells in culture retain markers of high glucose stress long after glucose levels have normalized (Ihnat et al., 2007).

H_2_S is common in the environment: in addition to natural sources of H_2_S, such as volcanic gasses and anaerobic bacteria in saline marshes, H_2_S is produced and emitted from large livestock farms, power plants, oil and natural gas refineries and pipelines, and during the production of glue, plastics, and asphalt (Beauchamp et al., 1984). The effects of longterm exposure to low H_2_S are poorly understood, but may contribute to neurological, respiratory, and cardiovascular dysfunction in humans (Kilburn and Warshaw, 1995; Richardson, 1995; Bates et al., 2002). However, exogenous H_2_S can also have beneficial effects in mammals, improving outcome in mammalian models of ischemia/reperfusion injury (Bos et al., 2015; Wu et al., 2015; Sen, 2017) and mediating at least some of the beneficial effects of dietary restriction (Hine et al., 2015).

*C. elegans* is an excellent model to understand the molecular and genetic pathways that mediate the responses to H_2_S and thereby the beneficial and/or detrimental physiological effects of H_2_S in animals. Just as exposure to high concentrations of H_2_S is lethal in mammals, *C. elegans* die when exposed to high concentrations of H_2_S (Budde and Roth, 2010). However, *C. elegans* grown in low H_2_S are long-lived and thermotolerant (Miller and Roth, 2007) and are better able to maintain proteostasis in hypoxia (Fawcett et al., 2015), suggesting that at least some protective effects of H_2_S observed in mammals may be recapitulated in the *C. elegans* model.

We have discovered a novel environmental bookmark in *C. elegans* formed by exposure to hydrogen sulfide (H_2_S). We show that transient exposure to low H_2_S leads to formation of a bookmark, which enables animals to survive subsequent exposure to otherwise lethal concentrations of H_2_S. This bookmark is quite stable, in that it can persist throughout development and far into adulthood. However, we have found that it is possible to reverse the bookmark by short periods of fasting. We further demonstrate that animals that have acquired the H_2_S bookmark induce the expression of some H_2_S-induced genes more robustly than naïve controls. These data suggest an epigenetic modification at the promoters of these bookmarked genes. Indeed, we demonstrate that mutations in the conserved histone H3 lysine 4 (H3K4) methyltransferase SET-2 and the H3K4me2 demethylase CoREST-like complex abrogate the ability to maintain the persistent effects of H_2_S exposure. Our results suggest that exposure to H_2_S results in changes in the methylation status of H3K4 at a subset of H_2_S-responsive promoters, which facilitates a robust transcriptional response that allows animals to survive subsequent H_2_S exposure.

## Results

### Exposure to low H_2_S forms an environmental bookmark

*C. elegans* grown in low H_2_S (50 ppm H_2_S in otherwise normal room air) have increased lifespan, increased thermotolerance, and improved maintenance of proteostasis (Miller and Roth, 2007; Fawcett et al., 2015). These animals raised in low H_2_S are also acclimatized, as they can survive when transferred to high H_2_S (concentrations higher than 150 ppm), a dose that is lethal to naïve controls (Budde and Roth, 2010). We repeated these experiments, confirming that animals grown in low H_2_S survived when transferred to high sulfide (Fig 1A). Although continuous exposure throughout development is required for H_2_S-induced thermotolerance and lifespan extension, shorter exposure to H_2_S is sufficient to protect against hypoxia-induced disruption of proteostasis (Miller and Roth, 2007; Fawcett et al., 2015). We therefore asked if shorter exposure to low H_2_S was sufficient to induce acclimatization. We exposed fourth-stage larvae (L4) *C. elegans* to low H_2_S for only 6 h and then immediately challenged animals with exposure to high H_2_S (150 ppm H_2_S in room air) overnight. Although all naïve animals exposed to high H_2_S died, 100% of animals that had a short acclimation to low H_2_S survived the subsequent exposure to high H_2_S (Fig 1A). This suggests that the physiological changes that occur in low H_2_S are relatively rapid and acclimatization does not require continuous growth in low H_2_S.

**Fig 1:**
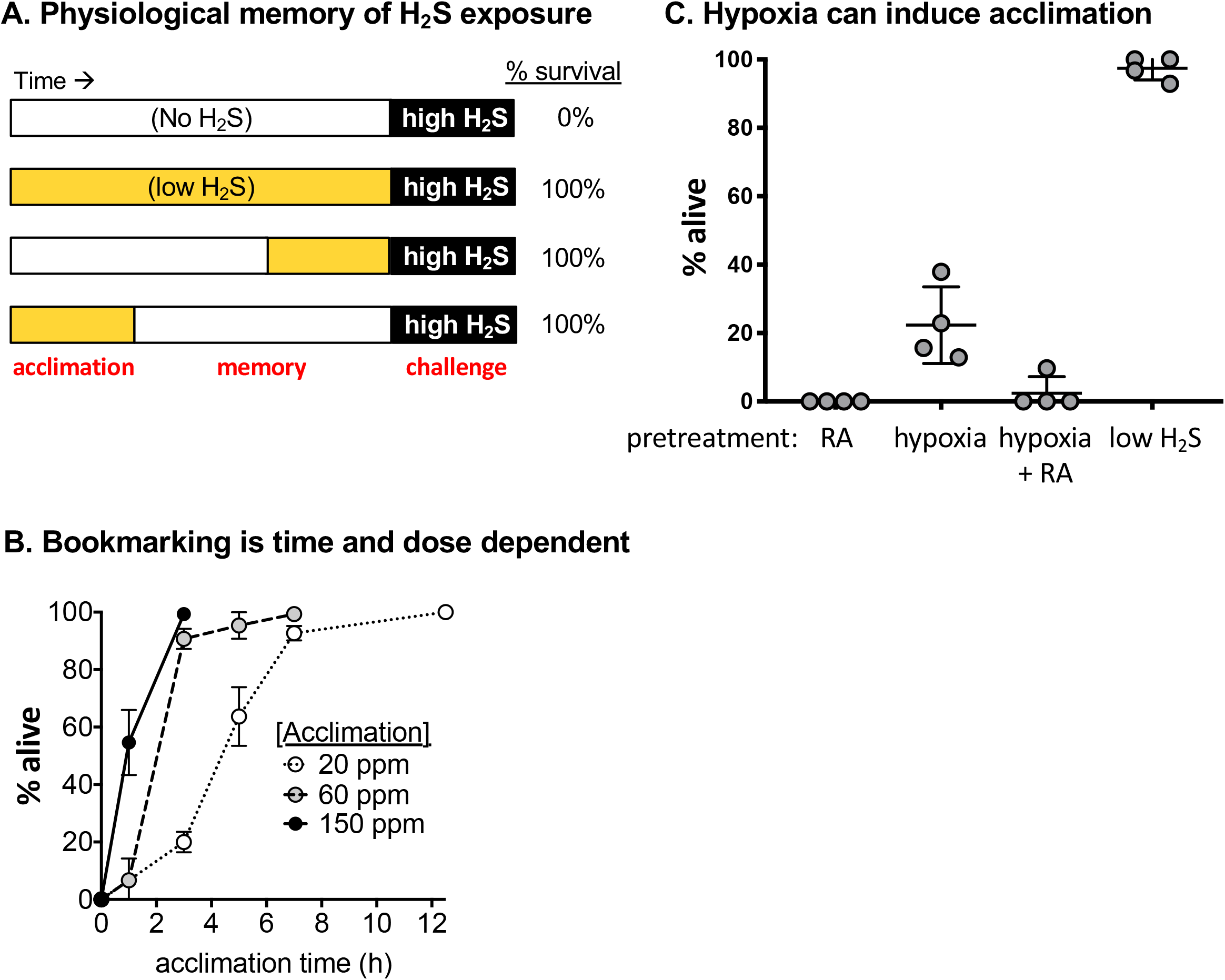
Exposure to low H_2_S induces an environmental bookmark. A. Survival of animals exposed to high H_2_S (black). In the schematic, time in low H_2_S is colored yellow, time in room air without H_2_S is white, and black indicates when animals were challenged with exposure to high H_2_S. TOP ROW: Naïve animals grown in the absence of H_2_S (open white). SECOND ROW, animals grown in low H_2_S (50 ppm) throughout development. THIRD ROW: Animals exposed to low H_2_S for 6 h as L4 and then immediately moved to high H_2_S. BOTTOM ROW: Embryos exposed to low H_2_S for 6 h, and then raised in room air without H_2_S for 48 h. As noted in red below the bottom row, we refer to the initial exposure to low H_2_S as acclimation, time in the absence of H_2_S as the memory period, and the subsequent exposure to high H_2_S as the challenge. Each exposure was repeated at least three times with at least 30 animals in each cohort. B. Time-dose response to form the H_2_S bookmark. Animals were acclimated to each indicated concentrations of H_2_S for the indicated time as LI, then challenged with 150 ppm H_2_S as L4. C. Survival of animals exposed to 150 ppm H_2_S after previous exposure to hypoxia. Wild-type L4 animals were exposed to 5,000 ppm 0_2_ for 5h and then either moved directly to high H_2_S (hypoxia) or allowed to recover in room air for 3h before exposure to high H_2_S (hypoxia + RA). Control animals remained in room air (RA) or were exposed to low H_2_S for 5 h (low H_2_S). Each point is the average survival from one independent experiment with at least 30 animals in each cohort for each experiment. Lines indicate the mean and standard deviation of the four independent experiments.

We next asked if animals that were acclimated to low H_2_S could form a persistent, physiological memory of the exposure to H_2_S. We exposed first-stage larvae (L1) to low H_2_S for 6h, then returned the animals to room air. When the animals reached L4 we challenged them with exposure to high H_2_S overnight. We found that all of the animals survived this treatment (Fig 1A). Thus, the effects of exposure to low H_2_S do not require direct transfer into high H_2_S, but instead can be maintained even in the absence of continued exposure to H_2_S. This suggests that there is a process that occurs in addition to acclimatization, which can be achieved simply by activation of HIF-1 transcriptional activity. We refer to this long-lasting physiological memory of H_2_S exposure as H_2_S bookmarking to distinguish it from acclimation.

The exposure to low H_2_S required to generate the H_2_S bookmark is time- and dose-dependent. We initially chose 50 ppm H_2_S for the low H_2_S exposure based on prior work demonstrating that this concentration of H_2_S causes clear physiological effects, such as increased lifespan and resistance to the hypoxia-induced disturbance of proteostasis (Miller and Roth, 2007, #51252; Fawcett et al., 2015). However, we found that animals exposed to only 20 ppm H_2_S developed the H_2_S bookmark and survived subsequent exposure to high H_2_S, though a longer exposure to the low H_2_S environment was required than at 50 ppm H_2_S (Fig 1B). Conversely, the bookmark was formed more rapidly at higher concentration of H_2_S, as even a 1h exposure to 150 ppm H_2_S was sufficient for survival of subsequent exposure. These data indicate that the bookmarking process can be quite rapid, but that the formation of the bookmark requires a threshold H_2_S exposure.

Stabilization of the HIF-1 transcription factor has been proposed to underlie the acclimatization to H_2_S (Budde and Roth, 2010). HIF-1 accumulates in animals exposed to H_2_S, and constitutive activation of HIF-1, from mutation of either *egl-9* or *vhl-1*, is sufficient for *C. elegans* to survive exposure to high H_2_S (Budde and Roth, 2010). We could not test directly whether *hif-1* is required for acclimatization or bookmarking, because *hif-1* mutant animals die when exposed to low H_2_S (Budde and Roth, 2010). However, we reasoned that if activation of HIF-1 was sufficient for acclimatization then animals exposed to low O_2_ (hypoxia) would also survive if transferred to high H_2_S. Consistent with this hypothesis, we observed increased survival of animals exposed to high sulfide when transferred directly from 5000 ppm O_2_, a condition where HIF-1 is activated (Fig 1C; Jiang et al., 2001, #24680). This result supports the hypothesis that activation of HIF-1 is sufficient for acclimation. However, the penetrance of survival was never as high as when animals were transferred from low H_2_S. This difference in penetrance could reflect the fact that although HIF-1 induces gene expression in both hypoxia and H_2_S, different transcripts accumulate in each condition (Miller et al., 2011). Alternatively, it could be that other factors that contribute to the process of H_2_S bookmarking are not engaged by exposure to hypoxia. We favor this model, because the acclimation induced by exposure to hypoxia is rapidly lost upon return to room air (Fig 1C). Thus, although the animals had acclimated they had not formed a persistent physiological memory consistent with H_2_S bookmarking.

We considered the possibility that the exposure to hypoxia activated stress response pathways that were cross-protective in H_2_S. Many well-studied stress response pathways are activated by and protective against multiple environmental stresses. For example, activation of the insulin-like signaling pathway leads to increased resistance to thermal, oxidative, and nutritional stress (McColl et al., 2010). Although exposure to H_2_S does not activate common stress-response pathways in *C. elegans*, including insulin-like signaling, TOR, and p53 signaling (Miller and Roth, 2009; Miller et al., 2011), we considered the possibility that protection by H_2_S bookmarking may be a result of the activation of such cross protective stress response pathways. We first tested whether inducing the heat shock response would protect against subsequent H_2_S exposure, as the heat-shock response is cross protective against several other stresses, including salt stress and oxidative stress, and animals grown in H_2_S are resistant to thermal stress (Völker et al., 1992; Miller and Roth, 2007; Verghese et al., 2012). We found that subjecting animals to a non-lethal heat shock did not protect against subsequent exposure to high H_2_S (Supplemental Fig 1A). Furthermore, animals with mutations in candidate cross-protective stress response signaling pathways, including those involved in the response to heat shock, nutrient deprivation, hypoxic stress, and osmotic stress did not have a defect in the formation or maintenance of H_2_S bookmarking (Supplemental Fig 1B). Taken together, these results suggest that H_2_S bookmarking is a specific response to H_2_S.

### H_2_S bookmarking is persistent, but reversible

During development, there are sometimes critical windows during which animals are most sensitive to certain environmental stimuli. For example, in *C. elegans*, the decision to enter dauer is determined by conditions during the first larval stage (Golden and Riddle, 1984). To determine if there was a specific developmental window in which H_2_S bookmarks could be formed, we exposed synchronized cohorts of *C. elegans* at different developmental stages to low H_2_S, and then tested their ability to survive exposure to high H_2_S after 48 h in room air. All animals survived subsequent exposure to high H_2_S, regardless of developmental stage at time of adaptation (Fig 2A). We conclude that H_2_S bookmarking can occur at any point in the lifetime of the animal.

**Fig 2:**
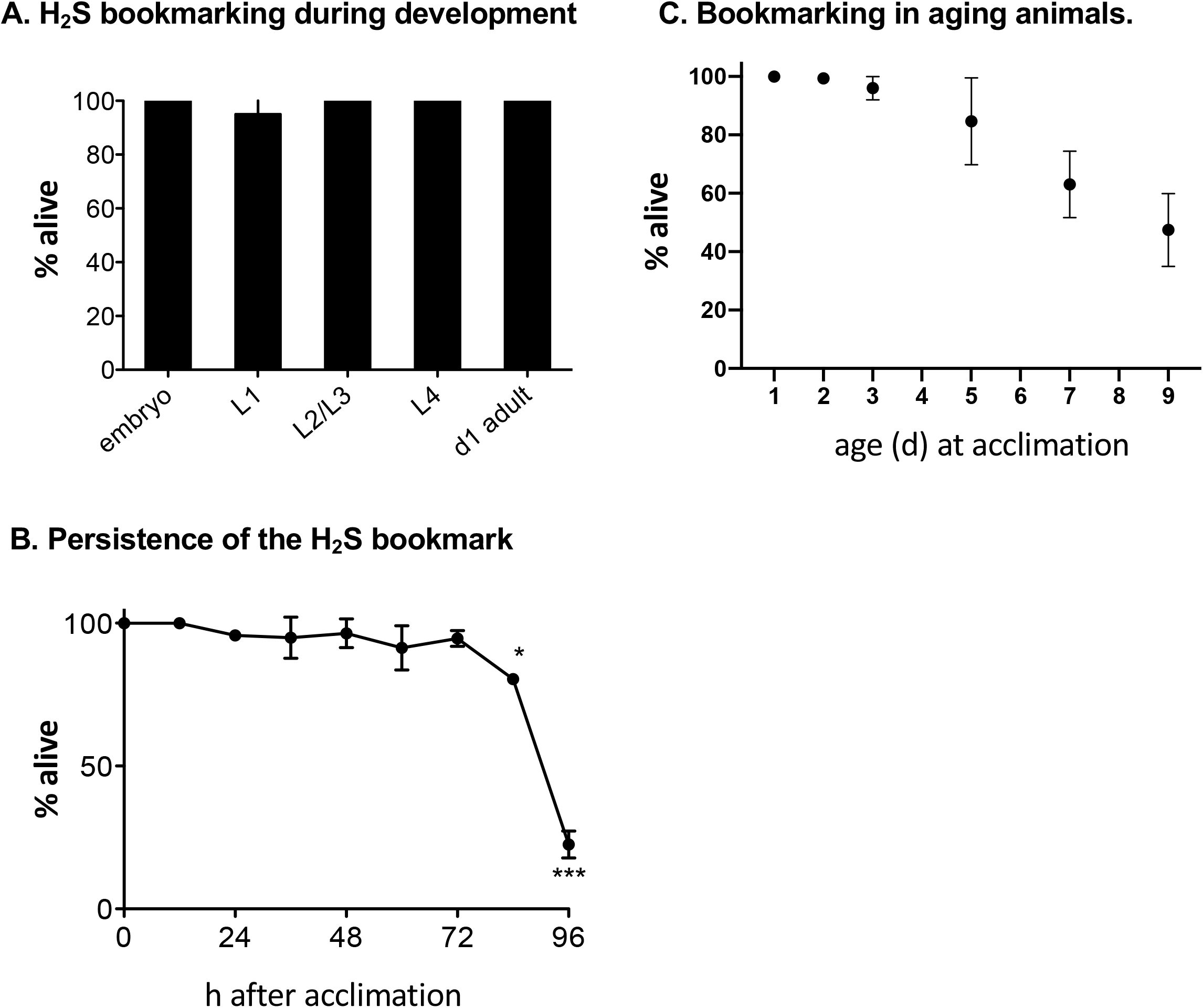
H_2_S bookmarks are persistent but reverse spontaneously with age. A. H_2_S bookmarking during development. Animals at each developmental stage were exposed to low H_2_S for 4 h, then returned to room air for 48 h before challenge with high H_2_S. B. H_2_S bookmark persistence. Animals were exposed to low H_2_S for 12 h as embryos and then returned to room air for the indicated period of time before challenge with high H_2_S. C. Bookmarking in aging animals. Animals were exposed to low H_2_S for 12 h at the indicated age, then returned to room air for 48 h before challenge with high H_2_S. For all panels, each experiment was repeated independently at least 4 times with 30-40 animals in each cohort. Error bars are standard deviation of the mean.

We next asked how long the H_2_S bookmark could persist. To measure this, we acclimated embryos to low H_2_S for 8 h, then let them grow in room air for increasing time before exposure to high H_2_S. We found that all animals exposed to H_2_S as embryos survived exposure to high H_2_S as much as 72h later, when the cohort were reproductive adults (Fig 2B). However, after 72h the survival of bookmarked animals began to decline and by 96h after the initial exposure most animals died when exposed to high H_2_S. The exact duration that the bookmark is maintained is somewhat dose-dependent, in that the decline began sooner if the acclimation period was shortened. These data indicate that the H_2_S bookmark can be maintained throughout embryonic and postembryonic development, but that it eventually spontaneously lost.

The fact that the bookmark spontaneously reversed as the animals aged led us to investigate the possibility that the rate of spontaneous reversal changed with age. To test this possibility, we exposed synchronized, aging cohorts of animals to low H_2_S for 6h and then let the animals age in room air for another 48h before challenging the animals with exposure to high H_2_S. We found that animals were able to form and maintain the H_2_S bookmark through day 3 of adulthood. However, starting at day 5 we observed a progressive decrease in the ability to survive the subsequent exposure to high H_2_S (Fig 2C). This result suggests that the H_2_S bookmark is formed or maintained less effectively in older animals. We did not observe a similar decline in acclimatization, as all animals survived when we transferred old animals to high H_2_S immediately after exposure to low H_2_S. Together these data suggest that H_2_S bookmarking – but not acclimatization – is less efficient with increasing age.

### H_2_S bookmarking is reversed by fasting

While characterizing the H_2_S bookmarking phenotype, we noticed that animals on plates in which the food source had been depleted no longer displayed robust survival in high H_2_S. We hypothesized that H_2_S bookmarking may be modulated by food availability. To test this possibility, we adapted embryos to low H_2_S for 12 hours on plates seeded with OP50 *E. coli* bacteria. We then removed half of the animals from food for 6-12 hours as first-stage larvae (L1), then returned these animals to normal food conditions. We challenged both fed and fasted animals with high H_2_S when they reached day 1 of adulthood. The proportion of animals that survived exposure to high H_2_S was significantly lower in the fasted animals (Fig 3A). This result suggests that fasting prevents maintenance of, or actively reverses, the H_2_S bookmark. This led us to next determine if fasting also prevented formation of the H_2_S bookmark. To test this possibility, we exposed starved L1 to low H_2_S for acclimation, then returned the animals to food in the absence of H_2_S. In contrast to animals acclimated to low H_2_S on food, animals exposed to low H_2_S in the absence of food did not survive subsequent exposure to high H_2_S (Fig 3B). These results suggest that both formation and maintenance of the H_2_S bookmark is inhibited by fasting.

**Fig 3:**
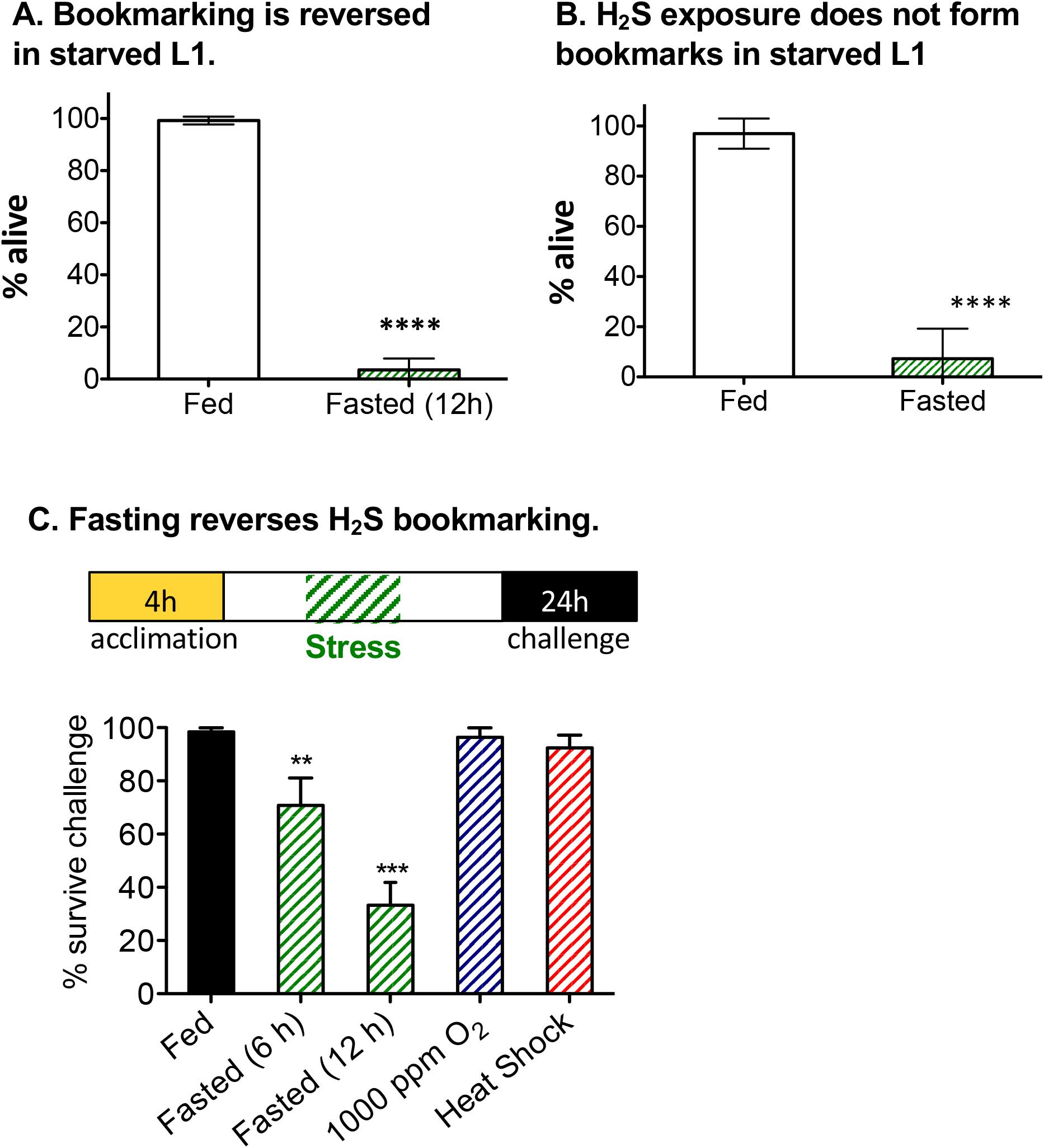
Fasting erases H_2_S bookmarks. A. Survival of animals exposed to high H_2_S when acclimated as embryos. Embryos were acclimated to low H_2_S for 4 h, then allowed to hatch in the presence or absence of food. After 12 h, L1 that hatched without food were moved to plates with food. When animals were L4 they were challenged with high H_2_S. B. Survival of animals acclimated to H_2_S in the absence of food. Embryos isolated from naïve adults were allowed to hatch in the absence of food overnight. Starved L1 were moved to plates ± food, then exposed to low H_2_S for 12 h. All animals were returned to house air and grown to L4/young adult on plates with food, then challenged with high H_2_S. C. H_2_S bookmarking is reversed by fasting, but not other stress responses. As indicated in the schematic, animals were exposed to low H_2_S for 4 h as embryos, then returned to room air. During the 48 h recovery time animals were transiently exposed to food deprivation (fasted; green), hypoxia (blue; 1,000 ppm O_2_ for 24h), or heat shock (red; 37 C for 1 h). The animals were then challenged with high H_2_S as L4/young adult. In all panels, graphs show mean survival ± standard deviation of at least 5 independent experiments with 30-40 animals in each experiment. ***, p < 0.001; **, p<0.01.

When *C. elegans* embryos hatch in the absence of food, starved L1 enter a reversible developmental arrest, the L1 diapause (reviewed in (Baugh, 2013)). To confirm that fasting itself reverses H_2_S bookmarking, and not the physiological changes associated with entry into the L1 diapause, we fasted animals at other developmental stages and scored for H_2_S bookmarking retention. We found that fasting at later developmental stages did erase the H_2_S bookmark (Fig 3C). Bookmarked animals removed from food for only 6 hours as L4s displayed a significant decrease in survival when later challenged with high H_2_S. This effect was more pronounced when the period of fasting increased. We conclude that fasting, or the response to food deprivation, effectively reverses the H_2_S bookmark.

One possibility is that activation of various stress responses, in addition to fasting, would erase the effect of the H_2_S bookmark. We therefore evaluated whether heat shock or hypoxia would reduce the ability of animals with an H_2_S bookmark to survive subsequent exposure to high H_2_S. We chose these two stresses because they both have global effects on physiology and metabolism, as fasting does. We found that neither hypoxia nor heat shock reversed the H_2_S bookmark (Fig 3C). Bookmarked animals exposed to hypoxia (1,000 ppm or 5,000 ppm O_2_; room air is 210,000 ppm oxygen) or heat shock survived exposure to high H_2_S as well as controls that remained in normal growth conditions. This result supports our assertion that the H_2_S bookmark is distinct from other general stress responses. Moreover, as hypoxia has dramatic effects to decrease protein translation and metabolic activity, this result suggests that maintenance of the H_2_S bookmark is not dependent on these processes.

### H_2_S bookmarking facilitates transcriptional reactivation of H_2_S responsive genes

One possible mechanism of H_2_S bookmarking could be that genes induced by exposure to low H_2_S persist, and are therefore able to promote survival in subsequent H_2_S exposure. We did not favor this hypothesis as the early transcriptional response to H_2_S depends entirely on *hif-1*, which is rapidly degraded in normal conditions (Wang et al., 1995; Kallio et al., 1999; Miller et al., 2011). To verify that H_2_S-induced changes in gene expression were not maintained in animals after exposure to low H_2_S, we first measured expression of the SQRD-1::GFP translational reporter, which is induced by exposure to low H_2_S (Budde and Roth, 2011). We exposed SQRD-1::GFP animals to low H_2_S for 8 hours, and then removed them to house air conditions for 48 hours. Expression of SQRD-1::GFP was significantly increased when animals were exposed to low H_2_S, but returned to baseline levels of expression upon return to house air (Fig 4A). To corroborate our results with SQRD-1::GFP, we measured the abundance of 12 H_2_S-induced transcripts (Miller et al., 2011). The abundance of these transcripts in bookmarked animals 48 h after exposure to low H_2_S was not significantly different from naïve controls (Fig 4B). We conclude that persistent changes in gene expression do not underlie the maintenance of the H_2_S bookmark.

**Fig 4:**
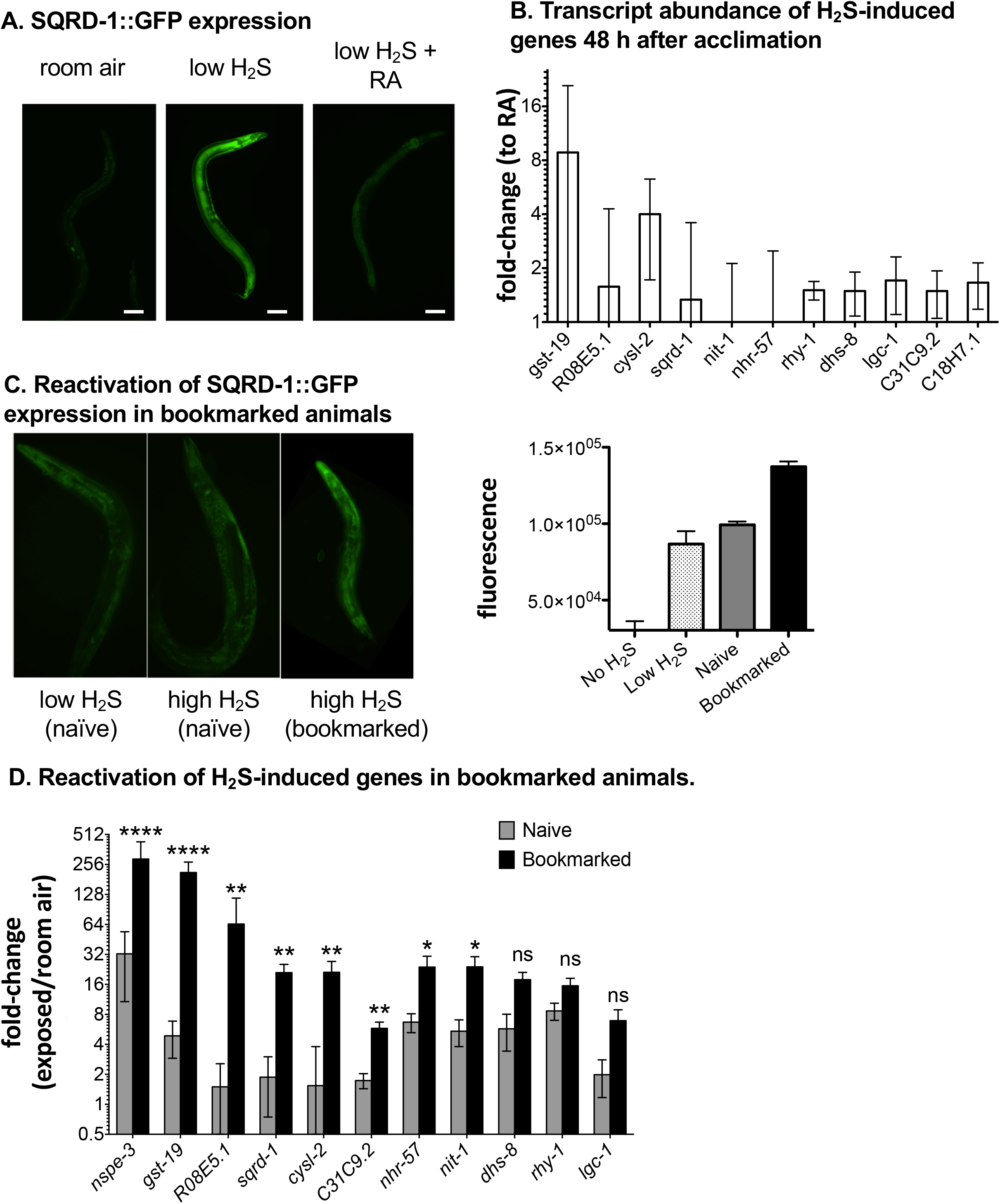
H_2_S bookmarking is associated with transcriptional reactivation of H_2_S-induced genes. A. Representative images showing expression of the SQRD-1::GFP translational fusion protein in the absence of H_2_S (room air; left), after 8 h in low H_2_S (middle), and then 48 h after return to room air (right). In all images anterior is toward the top. B. qRT-PCR measurement of abundance of mRNA from known H_2_S-induced genes. Animals were exposed to low H_2_S for 12 h and then returned to room air for 48 h. Graph shows mean transcript abundance relative to room air controls. Error bars are standard deviation of at least three experimental replicates. ns: not significant. C. Representative images showing expression of the SQRD-1::GFP translational reporter in naïve animals exposed to low or high H_2_S, and in bookmarked animals exposed to high H_2_S. All animals were first-day adults. Representative images are shown, with measured fluorescence intensity of cohorts of 30-40 animals shown in the graph. D. Transcriptional reactivation of H_2_S-inducible genes in bookmarked animals. Transcript abundance of known H_2_S-inducible genes was measured by qRT-PCR after animals were exposed to high H_2_S for 1 h. The change in transcript abundance relative to untreated controls is shown for naïve and bookmarked animals. Mean ± standard deviation is shown. ****, p-value < 0.0001; **, p-value < 0.01; *, p-value < 0.05; ns, not significant.

We next considered the possibility that the H_2_S bookmark functions to potentiate H_2_S-induced changes in gene expression upon subsequent re-exposure to H_2_S. We measured the expression of SQRD-1::GFP in naïve and bookmarked animals exposed to high H_2_S for 1 hour. We chose a 1-hour exposure because previous studies showed that expression of *sqrd-1* is significantly upregulated after 1 hour in low H_2_S (Miller et al., 2011), and naïve animals survive a 1-hour exposure to high H_2_S (Budde and Roth, 2011). We found that the increase in SQRD-1::GFP expression was similar when naïve animals were exposed to low or high H_2_S for 1 hour. In contrast, expression of SQRD-1::GFP was significantly higher when animals with the H_2_S bookmark were exposed to high H_2_S (Fig 4C). This suggests that the H_2_S bookmark facilitates the expression of H_2_S-inducible genes. To corroborate this result, we measured the abundance of transcripts upregulated by exposure to low H_2_S (Miller et al., 2011). Eight of the 12 genes we tested were increased more in bookmarked animals than naïve controls when exposed to high H_2_S (Fig 4D. We conclude that H_2_S bookmarking facilitates the transcriptional re-activation of H_2_S-responsive genes, which we refer to as bookmarked genes.

To get a more comprehensive understanding of how H_2_S bookmarking affected gene expression we performed RNAseq experiments to measure the abundance of transcripts in naïve and bookmarked animals exposed to high H_2_S. We found 16 transcripts that were upregulated in both bookmarked and naïve animals exposed to high H_2_S; of these, 12 were also identified in microarray experiments (Table 1; (Miller et al., 2011)). We also observed that 12 gene products were induced only in the naïve animals. None of these were upregulated in our previous microarray experiments. These genes could be induced specifically by exposure to high H_2_S, or it could be that they were false negatives (or missing) from the previous microarray experiment. Nevertheless, these studies suggest there is much in common between the transcriptional responses to low and high H_2_S.

**Table 1:**
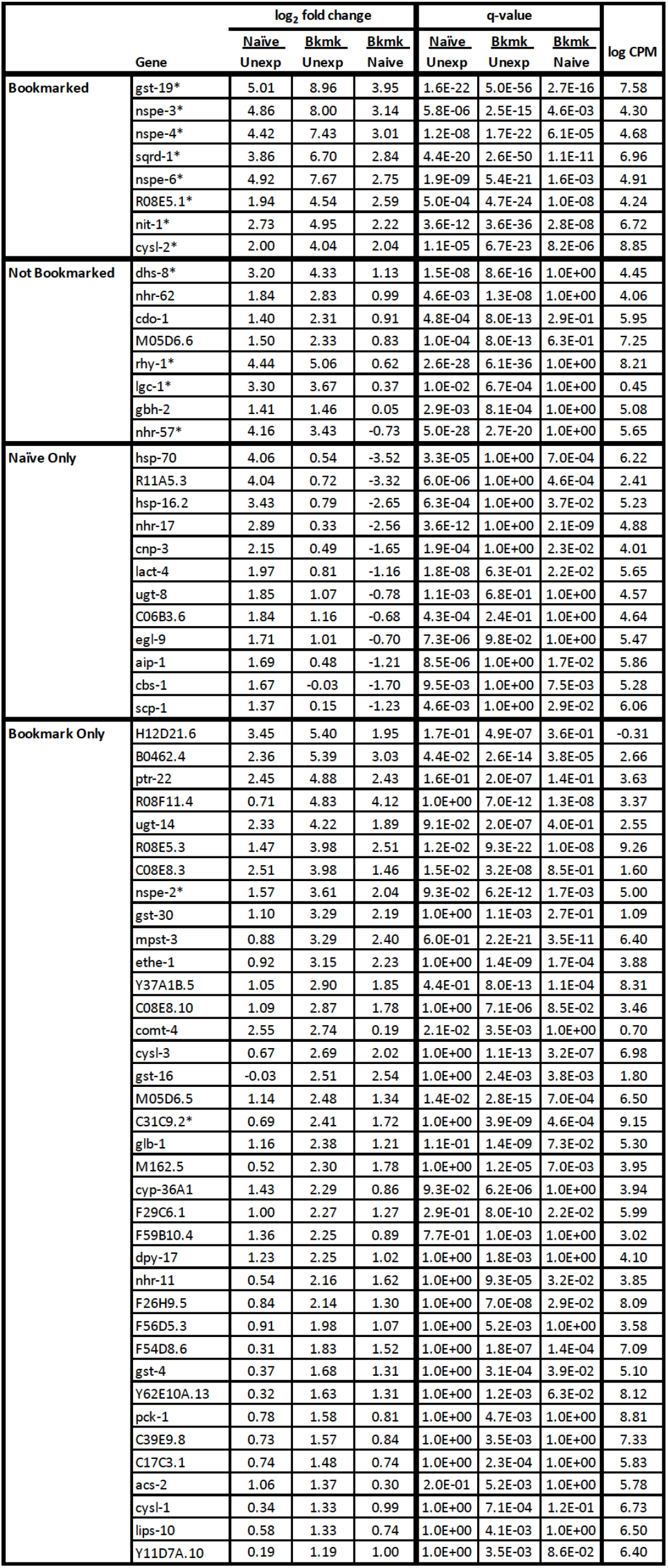
RNAseq comparison of gene expression in naïve and bookmarked animals. Differentially regulated genes after exposure to high H_2_S identified by RNAseq. Bookmarked genes were upregulated significantly more in animals previously exposed to low H_2_S than in naïve controls (but were induced by exposure to high H_2_S in the naïve animals). Not bookmarked genes were upregulated the same in treated animals and naïve controls. Naïve only genes upregulated only in the previously untreated controls. Bookmark only genes were significantly induced only in animals that had previously been exposed to low H_2_S.

We observed that eight transcripts were significantly more abundant in the bookmarked animals than in the naïve controls by RNA-seq (Bookmarked genes in Table 1). These results corroborated our qRT-PCR experiments (Fig 4D). In both experiments, gene products for*gst-19, nspe-3*, R08E5.1, *sqrd-1, cysl-2*, and *nit-1* were more abundant in bookmarked animals than in naïve controls. We also observed 37 transcripts that were induced by exposure to high H_2_S only in the bookmarked animals (Bookmark Only in Table 1). Most of these had not been previously shown to be induced by exposure to H_2_S. However, this group did include C31C9.2, which was bookmarked in our qRT-PCR experiments (Fig 4D), and *nspe-2*, which was upregulated in microarray experiments (Miller et al., 2011) but not included in our qRT-PCR panel. It may be that some of these genes are minimally induced by exposure to low H_2_S, and do not meet our cut-offs for determining significance. Together, our data suggest a model where H_2_S bookmarking leads to changes in transcriptional accessibility of specific H_2_S-inducible genes, which facilitates robust transcriptional reactivation upon subsequent H_2_S exposure.

We noted that not all H_2_S-responsive genes were bookmarked. In both RNAseq and qRT-PCR experiments expression of three gene products, *dhs-8, rhy-1*, and *lgc-1*, were induced the same in bookmarked animals and naïve controls exposed to high H_2_S. Several other genes in our RNAseq data were also induced equally in bookmarked and naïve animals. There is no pattern we could detect that predicted whether an H_2_S-responsive gene would be bookmarked for transcriptional reactivation.

### Epigenetic factors are required to maintain H_2_S bookmarks

We hypothesize that there are at least two processes required for H_2_S bookmarking: the original formation of the bookmark, which could be related to acclimation, and maintenance of the bookmark after the end of the exposure to H_2_S. We were interested in the mechanism by which the bookmark was maintained, as the rapid acquisition and reversibility of the H_2_S bookmark suggested that it may be mediated by an epigenetic mechanism (Mirbahai and Chipman, 2014). Alternatively, it is possible that bookmarking could depend upon the persistence of a long-lived protein or small RNA, or through another uncharacterized mechanism. To distinguish between these possibilities, we performed a candidate screen to identify molecular factors that mediate H_2_S bookmarking.

We screened animals with genetic mutations in machinery involved in protein turnover, RNA processing, epigenetic modifications, or factors associated with environmental bookmarks in other systems (a list of all 250 candidates tested is included as Supplemental Table 1). To identify mutations that specifically disrupted the maintenance of the H_2_S bookmark, we exposed synchronized embryos of each mutant strain to low H_2_S for 8 hours, returned them to house air for 48 hours, and then exposed them to high H_2_S for 24 h. We selected candidates that had reduced survival after exposure to high H_2_S, as compared to wild-type controls that acclimate and survive the subsequent exposure to high H_2_S. Two of the mutations that caught our attention in this screen were *spr-5* and *set-2* (Fig 5A). For the remainder of this work we focus on these two genes, as they regulate methylation of lysine 4 on histone 3 (H3K4me) and post-translational modifications of histone proteins occupying promoters can influence transcriptional levels (reviewed in Filipescu et al., 2014).

**Fig 5:**
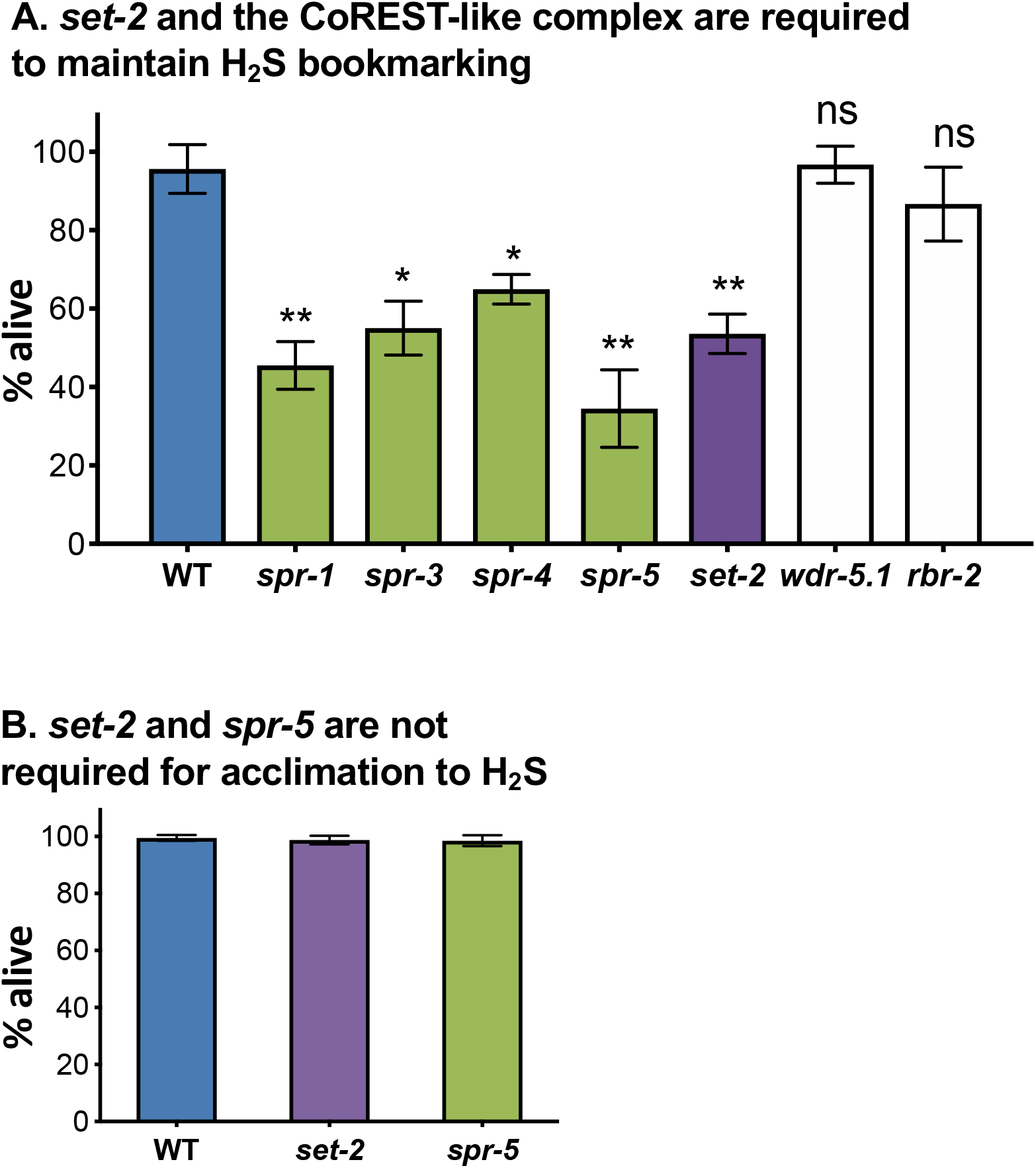
The SET-2 histone demethylase and CoREST histone methyltransferase are required to maintain H_2_S bookmarking. A. Bookmarking defects of *set-2* and CoREST-like complex members. Animals were exposed to low H_2_S for 4 h as embryos, then grown in room air for 48 h. Survival of challenge in high H_2_S is shown. The *set-2* (purple) and *spr-5* (green) mutant phenotypes were found in a candidate screen of 250 genes (listed in Supplementary Table 1). *spr-5, spr-1, spr-3*, and *spr-4* are members of the CoREST-like complex, *wdr-5.1* is a component of the ASH complex, and *rbr-2* is a histone demethylase that counteracts the ASH methyltransferase (see references in main text). Graph shows mean ± standard deviation of at least 5 independent experiments with 30-40 animals in each cohort. B. Acclimation of *set-2* and *spr-5* mutant animals. Animals were exposed to low H_2_S for 8h as L4 and then moved directly to high H_2_S overnight. Average survival is shown. Error bars are standard deviation of the mean. Graph shows mean ± standard deviation of at least 5 independent experiments with 30-40 animals in each cohort.

We were most interested in mutations that disrupted the long-term maintenance of the H_2_S bookmark, but we expected that some of the candidates identified in the primary screen could have defects in initial acclimation to H_2_S and/or formation of the H_2_S bookmark. To exclude that *spr-5* and *set-2* were required for acclimation, we exposed each candidate strain to low H_2_S for 6 hours, and immediately moved them to high H_2_S. We reasoned that if these animals were able to acclimate to H_2_S that they would survive this transition. Both *spr-5* and *spr-2* mutant animals survived this treatment (Fig 5B), suggesting that acclimation was not abrogated by these mutations.

H3K4 can be mono-, di, and tri-methylated. In *C. elegans*, the histone methyltransferase SET-2 is responsible for the formation of the majority of the bulk H3K4me2 and H3K4me3 (Xiao et al., 2011). However, formation of H3K4me3 by SET-2 requires other components of the ASH-2 methyltransferase complex. The ASH-2 complex in *C. elegans* is composed of three essential subunits: SET-2, ASH-2, and WDR-5.1 (Greer et al., 2010). To determine if SET-2 was acting as part of the ASH-2 complex in H_2_S bookmarking, we tested whether *wdr-5.1(ok1417)* mutant animals were capable of H_2_S bookmarking. In contrast to *set-2(n4589)* mutant animals, we found that *wdr-5.1(ok1417)* mutant animals survived exposure to high H_2_S after bookmarking at levels comparable to WT controls (Fig 5A). Similarly, mutations in *rbr-2*, an H3K4me3 demethylase that antagonizes ASH-2 methyltransferase complex activity (Greer et al., 2010, #35493), has no effect on H_2_S bookmarking (Fig 5A). Although SET-2 is unable to mediate formation of H3K4me3 independently of the ASH-2 complex, generation of H3K4me2 does not require the ASH-2 complex (Xiao et al., 2011). We therefore favor the hypothesis that SET-2 contributes to H_2_S bookmarking through its activity to form H3K4me2-modified histones.

SPR-5 is a component of the histone demethylase CoREST-like complex in *C. elegans* (Eimer et al., 2002). The CoREST complex was first identified in mammals as a corepressor of the REST transcription factor. Through its histone demethylase activity, the mammalian CoREST complex mediates long-term gene repression that is essential for the maintenance of cell identity (Andrés et al., 1999). In *C. elegans* the CoREST-like complex has retained its histone demethylase activity and functions in the repression of the *hop-1* gene through demethylation of H3K4me2 at the *hop-1* gene promoter. For *hop-1* gene repression, SPR-5 interacts with the orthologue of mammalian CoREST, SPR-1, and two large proteins with weak similarity to the mammalian REST transcription factor, SPR-3 and SPR-4 (Eimer et al., 2002). While they have no clear vertebrate homologs, SPR-3 and SPR-4 are predicted to function in the recruitment of the CoREST-like corepressor complex to gene targets. To determine if SPR-5 is functioning as a part of the CoREST-like complex in H_2_S bookmarking, we examined the effect of a loss-of-function mutation in these other known subunits of the CoREST-like complex. We found that, like *spr-5(by134)* mutant animals, animals with mutations in *spr-1, spr-3*, and *spr-4* all had defects in maintaining the H_2_S bookmark after exposure to low H_2_S (Fig 5A). Taken together, these results suggest that SPR-5 functions as a member of the CoREST-like complex to mediate H_2_S bookmarking.

We hypothesized that the transcriptional reactivation we observed in bookmarked animals was mediated by changes in H3K4me2 status at bookmarked promoters based on the genetic requirement for *set-2* and the CoREST demethylase complex. This hypothesis predicts that *set-2(n4589)* and *spr-5(by134)* mutant animals would not be able to effectively reactivate transcription of H_2_S-inducible transcripts. We tested this possibility by measuring transcript abundance in *set-2(n4589)* and *spr-5(by134)* mutant animals exposed to high H_2_S. Both *set-2(n4589)* and *spr-5(by134)* mutant animals exposed to low H_2_S as embryos had defects in the transcriptional reactivation of H_2_S-inducible genes (Fig 6A). One trivial explanation for the lack of transcriptional reactivation of H_2_S-inducible genes in the *set-2* and *spr-5* mutant animals is that they are generally unable to induce gene expression in response to H_2_S. We considered this unlikely, as both *set-2(n4589)* and *spr-5(by134)* mutant animals survive prolonged exposure to low H_2_S as well as wild-type animals. Indeed, we found no difference in the initial transcriptional response to low sulfide in either *set-2(n4589)* or *spr-5(by134)* mutant animals (Fig 6B). Moreover, both *set-2(n4589)* and *spr-5(by134)* mutant animals upregulated H_2_S-responsive genes as effectively as wild-type controls when exposed to high H_2_S after acclimation (Fig 6C). These results indicate that neither *set-2* nor *spr-5* are required for the normal transcriptional response or acclimatization to H_2_S. Instead, our data show that these histone-modifying enzymes are specifically required for the maintenance of the H_2_S bookmark and associated transcriptional reactivation of bookmarked genes.

**Fig 6:**
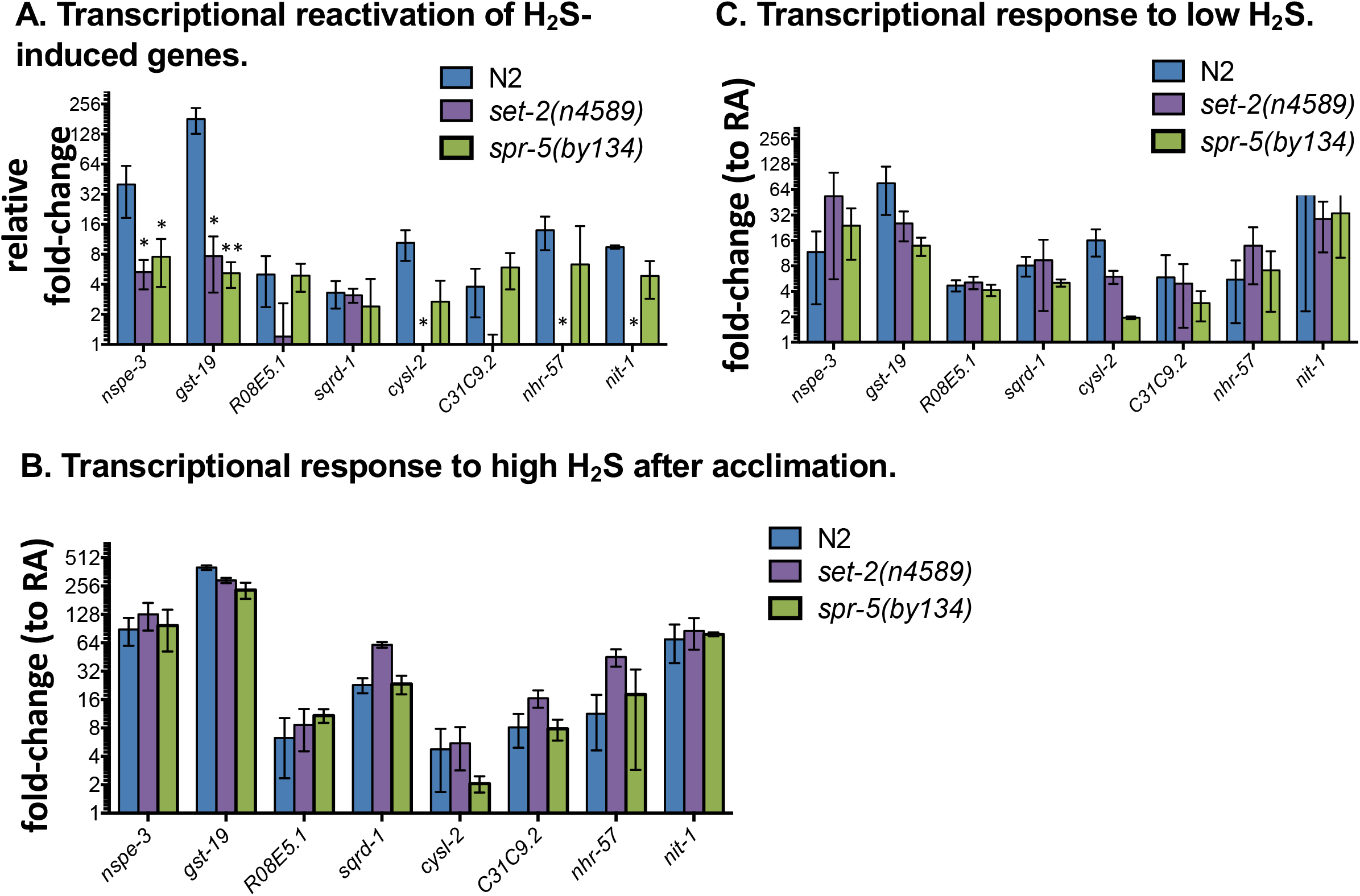
Transcriptional reactivation in bookmarked animals requires *set-2* and *spr-5*. A. Transcript levels were measured by qRT-PCR for naïve and bookmarked animals exposed to high H_2_S for 1 h. The relative fold-change of transcript abundance *(ÁAC_t_(bookmarked)/ AAC_t_(naïve))* is shown ± standard deviation. In all panels, N2 is in blue, *set-2(n4589)* is in purple, and *spr-5(by134)* is in green. In all panels ****, p-value < 0.0001; **, p-value < 0.01; *, p-value < 0.05; if not indicated, not significant. B. The initial transcriptional response to low H_2_S does not require *set-2* or *spr-5*. Transcript abundance for each H_2_S-inducible gene was measured in *set-2(n4589)* and *spr-5(by134)* mutant animals after 1 h exposure to low H_2_S. Graph shows mean change in transcript abundance, relative to untreated controls (ΔΔC_t_ ± standard deviation). C. *set-2* and *spr-5* mutant animals can be preconditioned to H_2_S. Animals of each genotype were exposed to low H_2_S for 24 h, then moved to high H_2_S for 1 h. The transcriptional response of these preconditioned animals is not distinguishable from wild-type controls. Graph shows ΔΔC_t_ ± standard deviation.

## Discussion

We have discovered a novel environmental bookmark to H_2_S that enhances survival when animals re-encounter H_2_S in the environment. Our experiments suggest that H_2_S exposure leads to formation of a bookmark that has persistent effects on the epigenetic landscape, allowing for enhanced transcriptional responses and increased survival upon subsequent H_2_S exposure.

To our knowledge, our studies demonstrate the first example of reversal of a stress-induced epigenetic bookmark by fasting. The molecular nature of the interaction between fasting and H_2_S bookmarking is not yet clear. In mammals, changes in diet, including periods of fasting, can lead to dramatic changes in the epigenome, including the reversal of epigenetic bookmarks (Burdge and Lillycrop, 2010). The ability to erase, or modulate, epigenetic bookmarks on demand has important implications in treatment of human disease by therapeutic intervention. Offspring of malnourished mothers can carry bookmarks that can lead to diabetes or obesity if food is abundant, which can reduce fertility (Gluckman et al., 2008). Maternal undernutrition during pregnancy, which results in epigenetic modifications to offspring that are designed to protect against subsequent famine, is associated with greater susceptibility to cancer in rats (Fernandez-Twinn et al., 2007). Our discovery of epigenetic bookmarking by H_2_S exposure provides a uniquely tractable model to develop strategies to modulate epigenetic marks in animals.

We propose that the H_2_S bookmark is related to the methylation status of histone H3K4, based on the genetic requirement for SET-2 and SPR-5 in maintaining the H_2_S bookmark. As SET-2 and SPR-5/CoREST both alter methylation status of H3K4me2, one simple model is that these two enzymes cooperate at the promoters of bookmarked animals to facilitate transcriptional reactivation. However, this interpretation is complicated by the fact that mutations in *set-2* and *spr-5* do not have entirely overlapping effects. For example, while transcriptional reactivation of*gst-19* and *nspe-3* was abrogated in both *spr-5* and *set-2* mutant animals, the bookmarking of *nhr-57, nit-1, C31C9.2*, and *R08E5.1* was lost only in *set-2* animals (Fig 5C). Moreover, some bookmarked genes were not affected in either *set-2(n4589)* or *spr-5(by134)* mutant animals. Future genome-wide studies of histone methylation status and the epigenetic effects of H_2_S are necessary to understand how these two histone-modifying enzymes are coordinated.

Our finding that H_2_S bookmarking requires histone methylation machinery is reminiscent of other established examples of epigenetic bookmarks. In plants, multiple rounds of drought stress lead to increased H3K4me3, providing resistance to subsequent severe droughts (Ding et al., 2012). Additionally, trithorax (TrxG) proteins with histone methyltransferase activity function in the epigenetic bookmarking of low winter temperatures in plants, which alters the timing of vernalization in subsequent seasons (Buzas et al., 2012; Song et al., 2012). The complexity and pleiotropic nature of environmental bookmarks has made understanding the mechanistic underpinnings of epigenetic bookmarks difficult, particularly in animals. Although not necessarily a bookmark, *C. elegans*, that enter into the alternative dauer larval stage have decreased in H3K4me3 and H4 acetylation in the adult animal, resulting in persistent changes in endo-siRNA levels that can poise genes for activation upon subsequent encounters with environmental stress (Hall et al., 2010).

The genetic factors required to maintain the H_2_S bookmark in *C. elegans*, SET-2 and SPR-5, are conserved through humans. Though to our knowledge H_2_S has not yet been shown to have epigenetic effects in mammals, H_2_S is a commonly encountered environmental toxin produced by oil refineries and paper mills that has dramatic effects on neurological, respiratory and cardiovascular function, even at low concentrations (Kilburn and Warshaw, 1995; Richardson, 1995; Bates et al., 2002). We demonstrate that H_2_S concentrations as low as 15 ppm, which is below OSHA limits for industrial exposure, can lead to the establishment of an H_2_S bookmark in *C. elegans*. It will be important to learn if H_2_S has similar effects in mammals, as this information could inform future toxic risk assessment to improve worker safety.

## Materials and Methods

*C. elegans* were maintained on nematode growth media (NGM) with OP50 *E. coli* at 20°C (Brenner, 1974). Worm strains used in this study are listed in Supplementary Table 1.

### Constructing defined gaseous environments

All compressed gas tanks were purchased from Airgas (Seattle, WA) and certified standard to 2% of target gas concentration, with the balance N2. Tanks with O_2_ contained 1,000 or 5,0 ppm, and H_2_S source tanks were 5,000 ppm. Gaseous environments were maintained using continuous flow chambers as described in (Fawcett et al., 2012). H_2_S was diluted from a 5,000 ppm stock tank with house air as in (Miller and Roth, 2007). H_2_S environments were maintained in a fume hood at 20°C, with matched house air (without H_2_S) continuous flow environments.

### H_2_S bookmarking assay

Embryos were synchronized by allowing gravid adults to lay eggs for 2 h on seeded NGM plates. First stage larvae (L1) were collected 24 h post egg-lay, L2 after 36 h, and L3s after 48 hours at 20°C on seeded NGM plates. L4 animals were picked from well-fed, logarithmically growing cultures and moved to seeded NGM plates. Staged animals were immediately exposed to 50 ppm H_2_S for indicated amount of time, and then removed to house air for 48 hours. Animals were then exposed to 150 ppm H_2_S overnight (~16 hours) and then scored for survival. Data was reported as *%* animals alive after 150 ppm H_2_S +/− standard deviation. Bagged animals were censored from the experiment. For differences between genotypes, p-values were calculated by one-way ANOVA using summary statistics (mean, standard deviation, n).

To evaluate the effects of other conditions on bookmarks, 24 h after the adaptation period animals were moved to hypoxia (1,000 or 5,000 ppm O_2_ for 24 h) or heat-shock (37 C for 1 h). For fasting, animals were moved to unseeded NGM plates with 25 mg/L carbenicillin to prevent bacterial growth. After 10 minutes, animals were moved to a new unseeded NGM plate with 25 mg/L carbenicillin to further deplete the food associated with their cuticle. Palmitic acid (10 mg mL^-1^ in ethanol) was used to form a physical barrier around the edge of each plate to encourage the animals to remain on the surface of the plate when fasted. After the indicated time period, animals were moved back to NGM plates seeded with live OP50 *E. coli* in room air until 48 h post-adaptation, at which time the animals were challenged with high H_2_S.

### qRT-PCR

Animals were grown on high-growth plates seeded with NA22 *E. coli* at 20°C. When animals reached gravid adult, synchronized embryos were obtained by a 5-minute bleach in 1:1:5 water:KOH:hypochloric acid solution. For each strain/condition, ~9,000 embryos were plated onto a 150 mM NGM plate seeded with live OP50 *E. coli*. Animals were not allowed to starve out the plate at any time during the experiment. The animals were treated as for the bookmarking assay as above, except that the exposure to 150 ppm H_2_S was for one hour. Animals were harvested into 1 mL Trizol solution and immediately frozen in liquid nitrogen. RNA was isolated from the Trizol preparation as described previously (Chomczynski, 1993). cDNA was made using Invitrogen SuperScript III First Strand Synthesis System. Primers were as in (Miller et al., 2011), primer sequences available upon request. qPCR was performed using Kappa SYBR FAST qPCR Kit. PCR cycle was as follows: 95C for 3 min, 95C for 15 sec, 55C for 15 sec x40. 4°C to hold. ΔC_t_ for each gene product was calculated as in (Miller et al., 2011); briefly, ΔC_t_ values were calculated by subtracting each measured C_t_ from the geometric mean of the control targets that are not altered in response to H_2_S (SIR-2.1, HIL-1, IRS-2, and TBA-1). ΔC_t_ were averaged across experiments. Student’s t-test was used to evaluate differences between ΔC_t_ values of treated samples and untreated controls. For differences between genotypes, p-values were calculated with a one-way ANOVA from summary statistics (mean, standard error, n). Reported fold-changes were calculated as 2^Λ^-ΔΔC_t_ where ΔΔC_t_ = ΔC_t_(experimental condition) – ΔC_t_(control condition). Error bars on graphs represent standard error of the mean, which was carried through the fold-change calculation using standard error propagation (www.statpages.org).

### GFP reporter quantification

Animals were synchronized by a 2-hour egg lay of SQRD-1::GFP animals onto seeded NGM plates. Animals were adapted to H_2_S as described above. After 48 hours, animals were exposed to 150 ppm H_2_S for 1 hour. Animals were then removed to house air and allowed to recover for 1 hour to allow for folding of GFP. Roller animals, which contain the SQRD-1::GFP transgene were mounted on an agar pad in a drop of 20 mM sodium azide as anesthetic. GFP fluorescence was visualized on a Nikon 90i fluorescent microscope with the GFP filter and oil-immersion 20x objective. All images were taken at the same exposure time and magnification. Total cell fluorescence was quantified using ImageJ software (Rasband, W.S., ImageJ, U. S. National Institutes of Health, Bethesda, Maryland, USA, http://imagej.nih.gov/ij/, 1997-2014). Student t-tests were used to compare mean cell fluorescence between samples.

### Global mRNA measurements with RNAseq

Embryos were synchronized by allowing gravid adults to lay eggs for 2-6 hours on seeded NGM plates. Within 24 hours the plates were exposed to 50ppm for 6 hours, and then removed to house air until animals reached L4, at which time they were exposed to 150ppm H_2_S for 1 hour, flash frozen in liquid nitrogen, and immediately stored at −80C.

RNA was isolated using Trizol, according to manufacturer’s instructions, and then isopropanol precipitation. Isolated RNA was sent to Novogene for library preparation using Illumina MiSeq for paired-end sequencing. The raw data files were uploaded to the Illumina analysis cloud (BaseSpace) in order to collect raw gene counts. Differential expression of genes between conditions was determined using the edgeR analysis package on the raw gene counts (Robinson et al., 2010).

## Acknowledgements

We thank our colleagues in the Miller lab that provided constructive comments to improve this manuscript and members of the NW Developmental Biology group that provided excellent feedback on this project. We are also thankful to Dr. R. Waterston (Department of Genome Sciences, University of Washington School of Medicine) for helpful advice and Dr. C. Chatterjee (Department of Chemistry, University of Washington) for insightful discussions. Mr. Harley Childs helped initiate the candidate screens for factors involved in H_2_S bookmarking. This work was supported by NIH grants R00 AG033055 and R01 ES024958 to DLM. DLM is an Ellison Medical Foundation New Scholar in Aging and has been supported by the Glenn Medical Foundation. EMF was supported by NIH Developmental Biology Predoctoral Training Grant T32 HD007183 from the NICHD. JJ was an Amgen Scholar. Some strains were provided by the CGC, which is funded by NIH Office of Research Infrastructure Programs (P40 OD010440).

## Author contributions

EMF performed experiments to characterize H_2_S bookmarking, identified epigenetic factors involved, and was involved in experimental design, data acquisition, analysis, and interpretation. EMF, CRB, and DLM wrote and edited early drafts of the manuscript. JKJ performed data acquisition and analysis, particularly in characterizing the effects of mutations in *set-2* and *spr-5*. EMG and CRB performed RNAseq experiments and analyzed data. DLM performed experiments in Fig. 1, planned experiments, and assisted in data analysis and interpretation. All authors edited and approved the final manuscript.

---

**Supplemental Figure 1.**
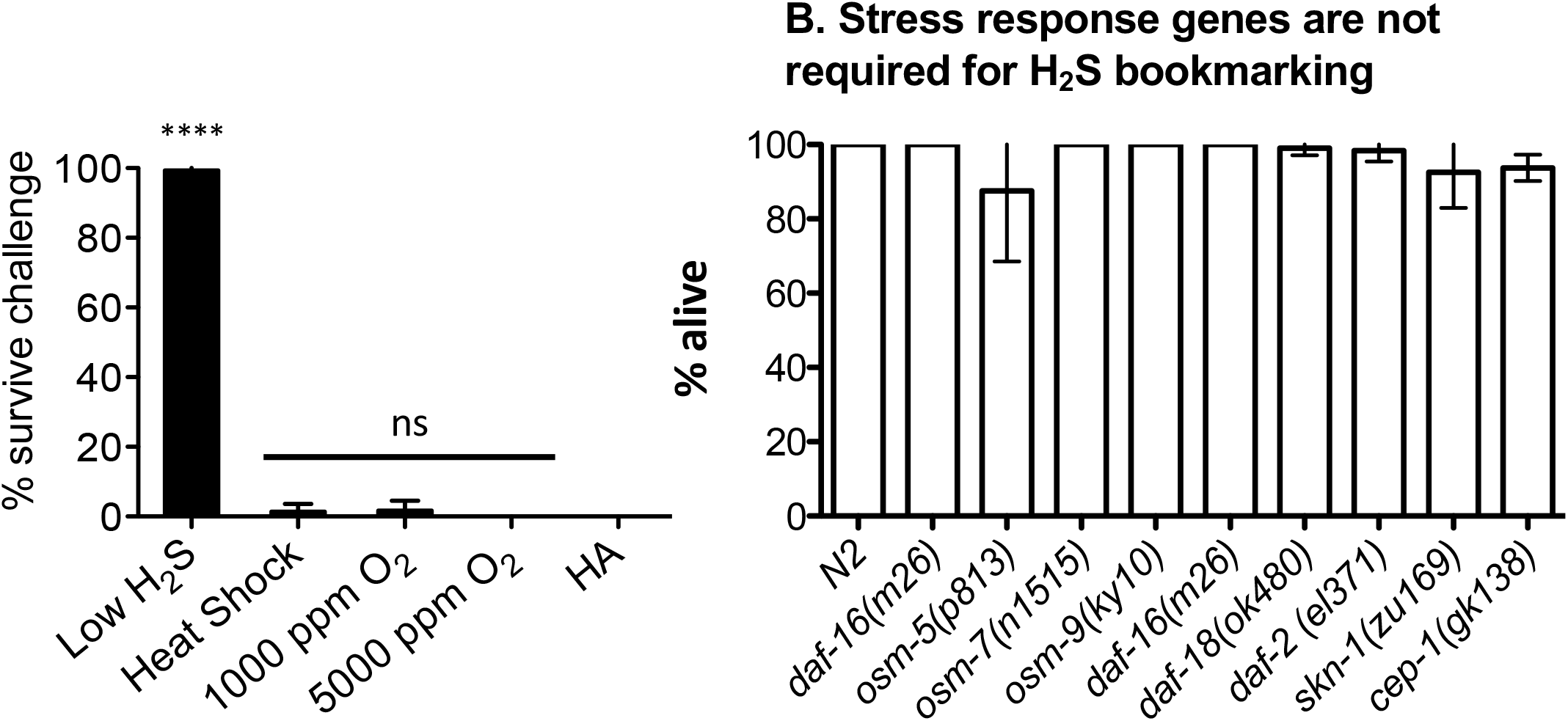
Bookmarking is specific to H_2_S. A. Survival of animals exposed to high H_2_S. L4 animals were exposed to either low H_2_S for 4 h, 37°C for 1 h (heat shock), or hypoxia (1,000 ppm or 5,000 ppm 0_2_) for 16 h. After treatment, animals were returned to room air. 48 hours later, animals were challenged with high H_2_S and scored for survival. B. General stress response pathways are not required for H_2_S bookmarking_ě_ Animals with mutations in genes involved in general stress responses were adapted to low H_2_S and challenged with high H_2_S 48 hours later. For both panels, each experiment was repeated at least five times, with 30-40 animals. Graphs shows mean +/− SD. Statistical comparisons were between controls (animals in house air (HA) or the N2 wild-type strain) and treatment groups: ****, p-value <0.0001; ns, not significant.

